# Theta Dual-Brain Stimulation of rTPJ Shapes Joint Agency

**DOI:** 10.64898/2026.03.10.710822

**Authors:** Yuto Kurihara, Ayaka Tsuchiya, Guillaume Dumas, Rieko Osu

## Abstract

Joint agency, the shared feeling of “we are doing this together”, has been linked to inter-brain synchrony, but its causal role in shaping this experience remains unclear. We applied dual transcranial alternating current stimulation (dual-tACS) over the right temporo-parietal junction (rTPJ) to 13 dyads performing an alternating tapping task (target ITI = 0.5 s; 180 deg. relative phase), manipulating in- and anti-phase coupling at theta (6 Hz), alpha (10 Hz), and beta (20 Hz). As a result, tapping in the theta anti-phase condition was significantly slower than the memorized reference tempo, whereas the other stimulation conditions did not influence the inter-tap interval. Meanwhile, the relative phase remained close to 180 deg. across all conditions. In the theta condition, anti-phase stimulation produced significantly lower joint agency than in-phase stimulation. Furthermore, mediation analysis suggested that the inter-tap interval may partially account for the effect of theta dual-brain stimulation on joint agency, although this indirect pathway did not reach statistical significance. These findings suggest that anti-phase theta stimulation over the rTPJ lowers joint agency, possibly by reducing coordination efficiency while preserving the overall 180 deg. alternation structure.

## Introduction

The subjective sense of causing or generating one’s own actions is a necessary component of self-consciousness. This phenomenon is defined as the sense of agency ^1–3^, the study of which has traditionally focused on individuals performing tasks alone ^4–6^. More recently, research on the sense of agency has expanded from individual to social contexts, particularly exploring the nature of this sense during joint action. Joint actions refer to activities where two or more agents coordinate their individual plans to collaboratively achieve a shared, external result ^7^. The sense of joint agency, described as the feeling that “we did it”, can emerge during coordinated group actions and differs from the sense of agency felt during individual actions ^8–10^. For example, Bolt et al. (2016) investigated the sense of joint agency by instructing pairs of participants to continuously coordinate their actions during a joint tapping task, showing that cooperative pairs tended to experience joint rather than self-agency, with this feeling becoming stronger as their mutual coordination increased ^11^. In addition, recent work has shown that joint agency can be stronger in musical duets than in constant-pitch synchronization, even when partners achieve comparable levels of interpersonal synchrony, suggesting that relational structure between partners’ actions plays a key role in shaping the experience of acting together ^12^. Taken together, studies on agency in joint action indicate that individuals can experience a sense of joint agency, and that this experience is shaped by available perceptual cues as well as the role they assume within the joint activity.

The neural mechanisms underlying the sense of joint agency remain unclear. Only a limited number of studies have directly examined how inter-brain synchrony between partners relates to their experience of joint agency. Inter-brain synchrony—defined as the temporal alignment of oscillatory activity across individuals—has been widely reported in hyperscanning studies of cooperative behavior, with evidence showing increased inter-brain synchrony in specific frequency bands (especially electroencephalogram (EEG) or magnetoencephalography (MEG)) and regions during social interaction ^13–15^. Within this literature, Shiraishi and Shimada (2021) demonstrated that theta-band synchrony between the prefrontal cortex and the right temporo-parietal junction (rTPJ) not only distinguishes cooperative from independent actions but also scales with the dyad’s subjective sense of joint agency ^16^. However, the causal relationship between inter-brain synchrony and joint agency remains unclear. It is an open question whether inter-brain synchrony is merely an epiphenomenon, arising from individuals independently processing common external stimuli, or if it is an emergent property that causally underpins their mutual coordination.

To test whether inter-brain synchrony causally enhances the sense of joint agency, we adopted a dual-brain stimulation protocol ^17–20^ during an alternating tapping task ^21,22^ (Figure 1A). This approach delivers transcranial electrical stimulation (tES) to two individuals at the same time (Figure 1B). By adjusting the coupling between the stimulation signals applied to each brain, experimenters can modulate inter-brain synchrony and assess its causal influence on social behavior ^18,23^.

**Figure 1.**
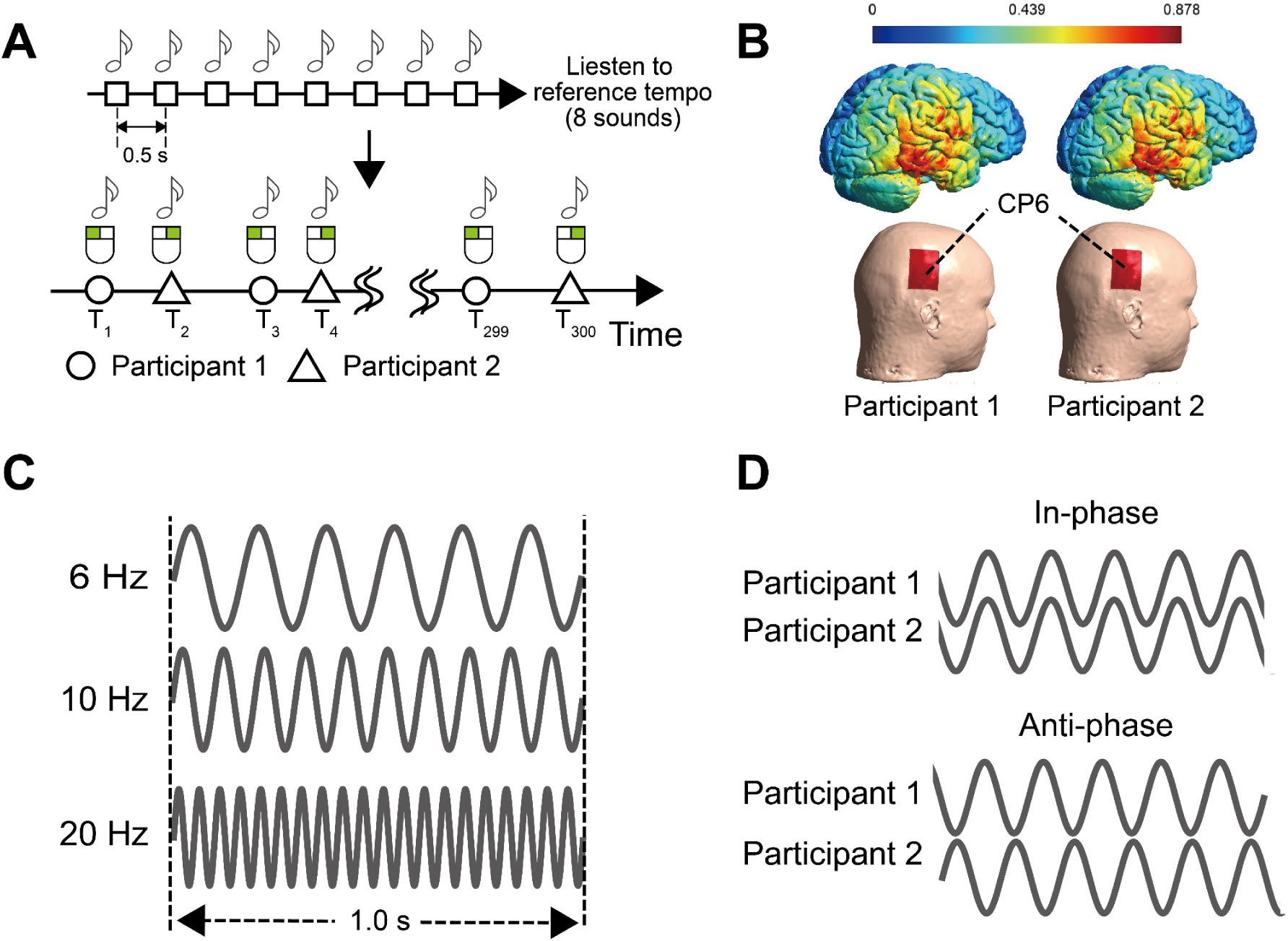
Experimental task and design. (A) Participant dyads performed an alternating tapping task using their right index fingers. This task was conducted over 6 trials. (B) Dual-tACS was applied during the alternating tapping task over electrodes placed at CP6 (anode) and contralateral forearm (cathode) to optimally target the right temporo-parietal junction. (C, D) The relative stimulation phase between brains was manipulated to be either in-phase (0° relative phase) or anti-phase (180° relative phase), across three stimulation frequencies: 6 Hz (theta band), 10 Hz (alpha band) or 20 Hz (beta band).

To investigate this causal link, previous studies have primarily utilized transcranial alternating current stimulation (tACS). tACS is a non-invasive method capable of entraining neural oscillations ^24^ with precise control over stimulation frequency ^25^ and phase ^26^. By applying phase-coupled tACS, researchers have demonstrated that inter-brain synchrony modulates specific behavioral outcomes depending on the targeted cortical region and oscillation frequency. For instance, Novembre et al. (2017) showed that in-phase 20 Hz (beta) tACS over the primary motor cortex enhances interpersonal movement synchrony ^18^. Similarly, targeting frontal regions such as the inferior frontal gyrus with 6 Hz (theta) or 20 Hz (beta) oscillations has been shown to facilitate social coordination and interactive learning ^17,19^. However, the effects are highly specific to the stimulation parameters. Szymanski et al. (2017) targeted right fronto-parietal regions with 6 Hz (theta) stimulation but did not observe enhancement in dyadic drumming performance ^20^. While these tACS studies successfully modulated behavioral performance (e.g., motor synchrony or learning accuracy), the present study aims to clarify the causal role of inter-brain synchrony in the subjective experience of joint agency. To this end, we employed the dual-tACS system to causally manipulate inter-brain synchrony and examine its effect on the sense of joint agency during the task.

Here, we targeted the rTPJs of each member of the dyad, simultaneously applying tACS to both participants at 6, 10, or 20 Hz (Figure 1C) in either an in-phase or anti-phase configuration (Figure 1D). The TPJ is widely regarded as a core hub for theory of mind, underpinning the ability to infer and represent others’ mental states ^4,27–29^. Previous hyperscanning studies have further reported synchronized activity in the TPJ during social interaction, particularly within the right hemisphere ^16,30–33^. Accordingly, we selected the rTPJ as the stimulation target for dual-tACS. The stimulation frequencies (6, 10, and 20 Hz) and phase configurations (in-phase and anti-phase) followed the protocol of Pan et al. (2021) ^19^.

Shiraishi and Shimada (2021) showed that inter-brain synchrony in the theta band is associated with the sense of joint agency ^16^. Based on this finding, we predicted that 6 Hz stimulation, which corresponds to the theta frequency range, would modulate joint agency in the present study. Pan et al. (2021) also demonstrated through mediation analysis that interpersonal movement synchrony partially mediates the effect of dual-brain stimulation on social outcomes. Based on this evidence, we further predicted that tapping-related variables would mediate the effect of dual-brain stimulation on joint agency. To assess tapping performance, we calculated the mean inter-tap interval (ITI), the mean relative phase (RP), and the standard deviation of RP (SDφ). The mean ITI reflects tapping speed (shorter ITIs indicate faster tapping), the mean RP captures the average phase relationship between the two tap trains, and SDφ quantifies variability in relative phase, providing an index of tapping instability (larger SDφ indicates less stable coordination). The subjective sense of joint agency was measured using visual analog scale (VAS) ratings. In addition, we used the Inclusion of Other in the Self (IOS) scale ^34^ to examine whether dual-tACS induced changes in perceived interpersonal closeness.

## Results

### Behavioral analysis

Behavioral analyses were conducted at the dyad level (13 dyads). To test whether mean ITI differed from the reference ITI (0.50 s), we used one-sample t-tests against 0.50 s for each condition. Mean ITI in the theta anti-phase condition was significantly longer than the reference ITI (t(12) = 4.65, padj = 0.0034, d = 1.29; Figure 2A). In contrast, the remaining five conditions—theta in-phase, alpha in-phase, alpha anti-phase, beta in-phase, and beta anti-phase—did not differ significantly from the reference ITI of 0.50 s, as indicated by one-sample t-tests (theta in-phase, t(12) = 0.609, p_adj_ = 0.554, d = 0.169; alpha in-phase, t(12) = 1.63, p_adj_ = 0.155, d = 0.45; alpha anti-phase, t(12) = 2.60, p_adj_ =0.069, d = 0.72; beta in-phase, t(12) = 2.10, p_adj_ = 0.113, d = 0.58; beta anti-phase, t(12) = 1.95, p_adj_ = 0.113, d = 0.54) (Figure 2A-C). We also performed a two-way repeated-measures ANOVA on mean ITI with factors Frequency (theta, alpha, beta) and Phase (in-phase, anti-phase). This analysis showed no significant Frequency × Phase interaction and no significant main effects of Frequency or Phase (Interaction, F(2, 24) = 2.703, p = 0.088, ηg² = 0.0221; Frequency, F(2, 24) = 0.030, p = 0.970, ηg² = 0.0003; Phase, F(1, 12) = 1.243, p = 0.287, ηg² = 0.0147). Taken together, these results suggest that theta anti-phase stimulation may increase the ITI relative to the target tempo, despite the absence of reliable between-condition differences in the ANOVA.

**Figure 2.**
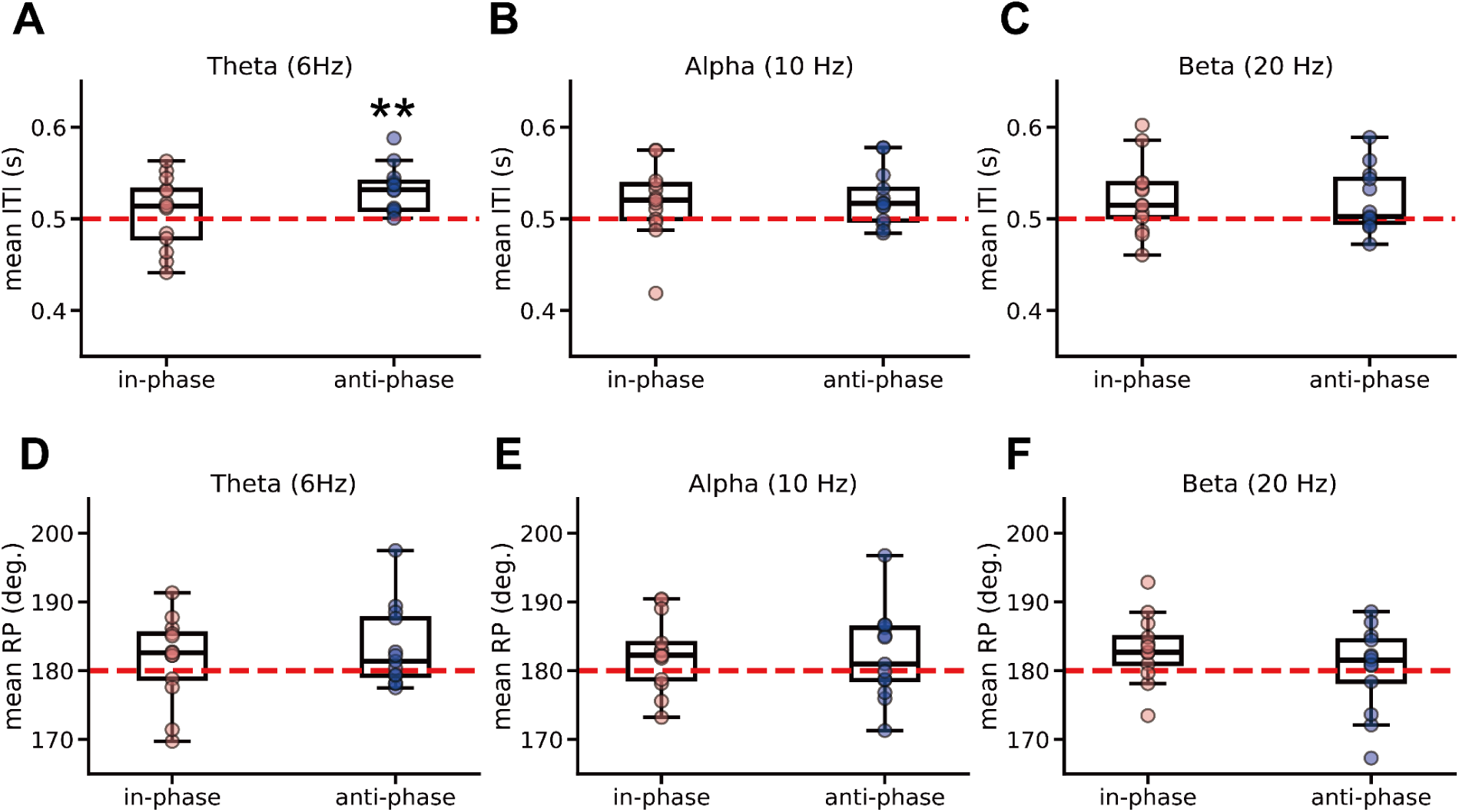
The results of inter-tap interval (ITI) and relative phase (RP). (A) Mean ITI under theta (6 Hz) anti-phase stimulation was significantly different from the 0.50-s reference, whereas no such difference was observed in the theta in-phase condition. (B) Mean ITI under alpha (10 Hz) in- and anti-phase stimulation did not differ significantly from 0.50 s. (C) Mean ITI under beta (20 Hz) in- and anti-phase stimulation did not differ significantly from 0.50 s. (D-F) In the theta, alpha, and beta bands, mean RP did not differ significantly from 180° in either the in-phase or anti-phase condition. **: p < 0.01

To assess whether mean RP deviated from the reference value of 180°, we conducted one-sample t-tests with μ = 180° for each condition. As a result, there were no significant differences from 180° for any condition (theta in-phase, t(12) = 1.05, p_adj_ =0.376, d = 0.29; theta anti-phase, t(12) = 1.86, p_adj_ =0.236, d = 0.72; alpha in-phase, t(12) = 1.57, p_adj_ =0.236, d = 0.44; alpha anti-phase, t(12) = 1.51, p_adj_ =0.236, d = 0.42; beta in-phase, t(12) = 1.75, p_adj_ =0.236, d = 0.48; beta anti-phase, t(12) = 0.056, p_adj_ =0.956, d = 0.016) (Figure 2D-F). A two-way repeated-measures ANOVA with factors Frequency and Phase on mean RP also showed no significant interaction or main effects (Interaction, F(2, 24) = 1.392, p = 0.268, ηg² = 0.022; Frequency, F(2, 24) = 1.398, p = 0.267, ηg² = 0.022; Phase, F(1, 12) = 0.0132, p = 0.910, ηg² = 0.00008). Collectively, these results indicate that tapping remained clustered around 180° across conditions, with no reliable differences in mean RP between phases or frequencies.

Furthermore, we calculated the standard deviation of RP (SDφ) to check the stability of their alternating tapping. A two-way repeated-measures ANOVA with factors Frequency and Phase on SDφ showed no significant interaction or main effects (Interaction, F(2, 24) = 1.047, p = 0.366, ηg² = 0.026; Frequency, F(2, 24) = 0.722, p = 0.496, ηg² = 0.0038; Phase, F(1, 12) = 1.821, p = 0.202, ηg² = 0.012) (Figure 3).

**Figure 3.**
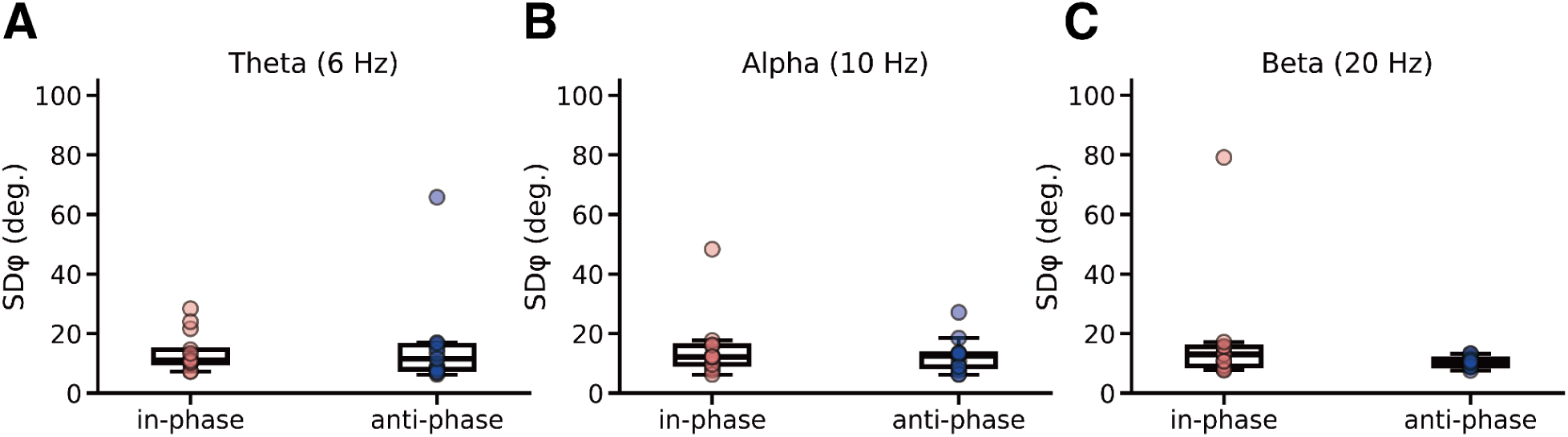
Results of SDφ. SDφ was defined as the circular standard deviation of the relative phase (RP). Across the combined conditions of Frequency (theta (A), alpha (B), beta (C)) and Phase (in-phase, anti-phase), no significant differences in SDφ were observed.

### Subjective scores analysis

Subjective ratings were analyzed at the individual-participant level (26 participants). We conducted two-way repeated-measures ANOVAs with factors Frequency and Phase for joint agency and IOS (Figure 4). For joint agency, the Frequency (6Hz, 10Hz, and 20Hz) × Phase (in-phase and anti-phase) interaction was significant (F(2, 50) = 4.052, p = 0.023, ηg² = 0.027), whereas neither factor showed significant main effects (Frequency, F(2, 50) = 2.910, p = 0.063, ηg² = 0.014; Phase, F(1, 25) = 0.154, p = 0.698, ηg² = 0.00058).

**Figure 4.**
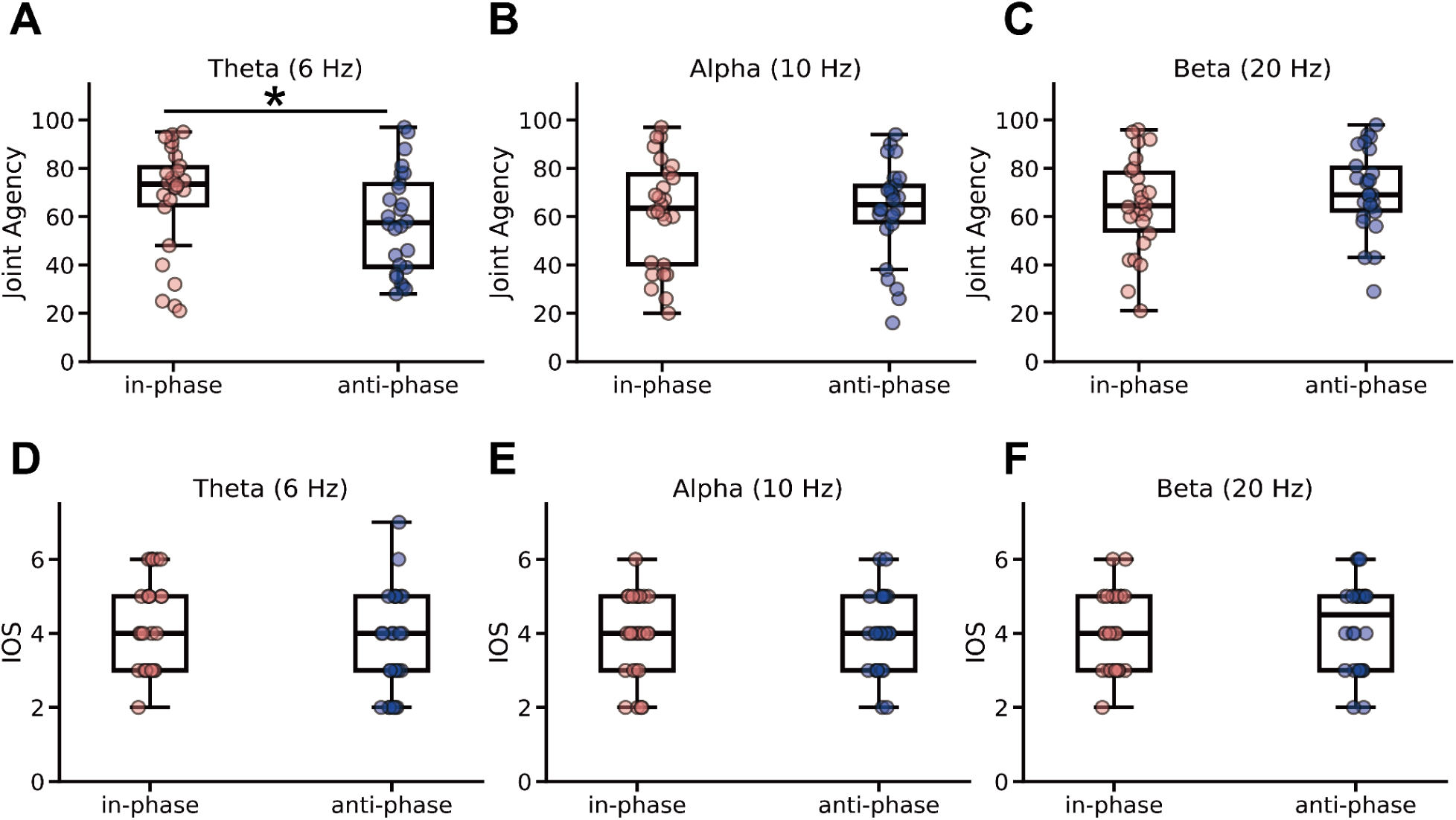
The results of the joint agency and Inclusion of Others in Self (IOS) scale. For joint agency, a two-way ANOVA with factors Frequency (theta, alpha, beta) and Phase (in-phase, anti-phase) revealed a significant Frequency × Phase interaction. Follow-up tests showed a significant Phase effect at 6 Hz (theta) (A), with in-phase ratings higher than anti-phase, whereas no Phase differences were observed at 10 Hz (alpha) (B) or 20 Hz (beta) (C). For IOS, a two-way ANOVA with factors Frequency (theta (D), alpha (E), beta (F)) and Phase (in-phase, anti-phase) revealed no significant Frequency × Phase interaction, and neither the main effect of Frequency nor the main effect of Phase was significant. *: p < 0.05

Follow-up simple-effects analyses examined the effect of Phase (in-phase vs. anti-phase) at each frequency. The effect of Phase was significant at 6 Hz (theta), F(1, 25) = 5.23, p = 0.031, ηg² = 0.052, indicating higher joint agency ratings in the in-phase than in the anti-phase condition. No significant Phase effect was observed at 10 Hz (alpha), F(1, 25) = 0.05, p = 0.83, ηg² = 0.001, or at 20 Hz (beta), F(1, 25) = 3.54, p = 0.072, ηg² = 0.027, though the latter showed a trend toward higher ratings in the anti-phase condition.

The simple main effect of Frequency was significant under the anti-phase condition (F(2, 50) = 4.851, p = 0.012, ηg² = 0.070) but not under the in-phase condition (F(2, 50) = 1.696, p = 0.194, ηg² = 0.014). Post-hoc pairwise comparisons (FDR-corrected) revealed that, under the anti-phase condition, joint agency ratings were significantly higher in the beta (20 Hz) than in the theta (6 Hz) condition (t(25) = 2.992, p_adj_ = 0.018, d = 0.66). The differences between beta and alpha (t(25) = -1.925, p_adj_ = 0.098, d = -0.429) and between alpha and theta (t(25) = 1.191, p_adj_ = 0.245, d = 0.231) did not reach significance.

A two-way repeated-measures ANOVA with factors Frequency (6, 10, and 20 Hz) and Phase (in-phase, anti-phase) was conducted on the IOS ratings (Figure 4D-F). Neither the main effect of Frequency ( *F*(2, 50) = 0.095, *p* = 0.910, ηg² = 0.001), nor Phase ( *F*(1, 25) = 2.990, *p* = 0.096, ηg² = 0.007), was significant. The interaction between Phase and Frequency was also not significant (*F*(2, 50) = 0.843, *p* = 0.436, ηg² = .005).

### Correlation and mediation analysis

Because only theta stimulation affected both tapping tempo and joint agency, subsequent analyses focused on this frequency. The correlation between raw ITI and joint agency in the theta condition is shown in Figure 5A. ITI and joint agency were negatively correlated in the in-phase condition (r = -0.398, p = 0.044), whereas no significant correlation was observed in the anti-phase condition (r = 0.146, p = 0.476). For the mediation analysis, ITI was log-transformed to reduce skew (Figure 5B). These results suggest that, in the in-phase condition, faster tapping (i.e., shorter inter-tap intervals) was associated with a greater sense of joint agency.

**Figure 5.**
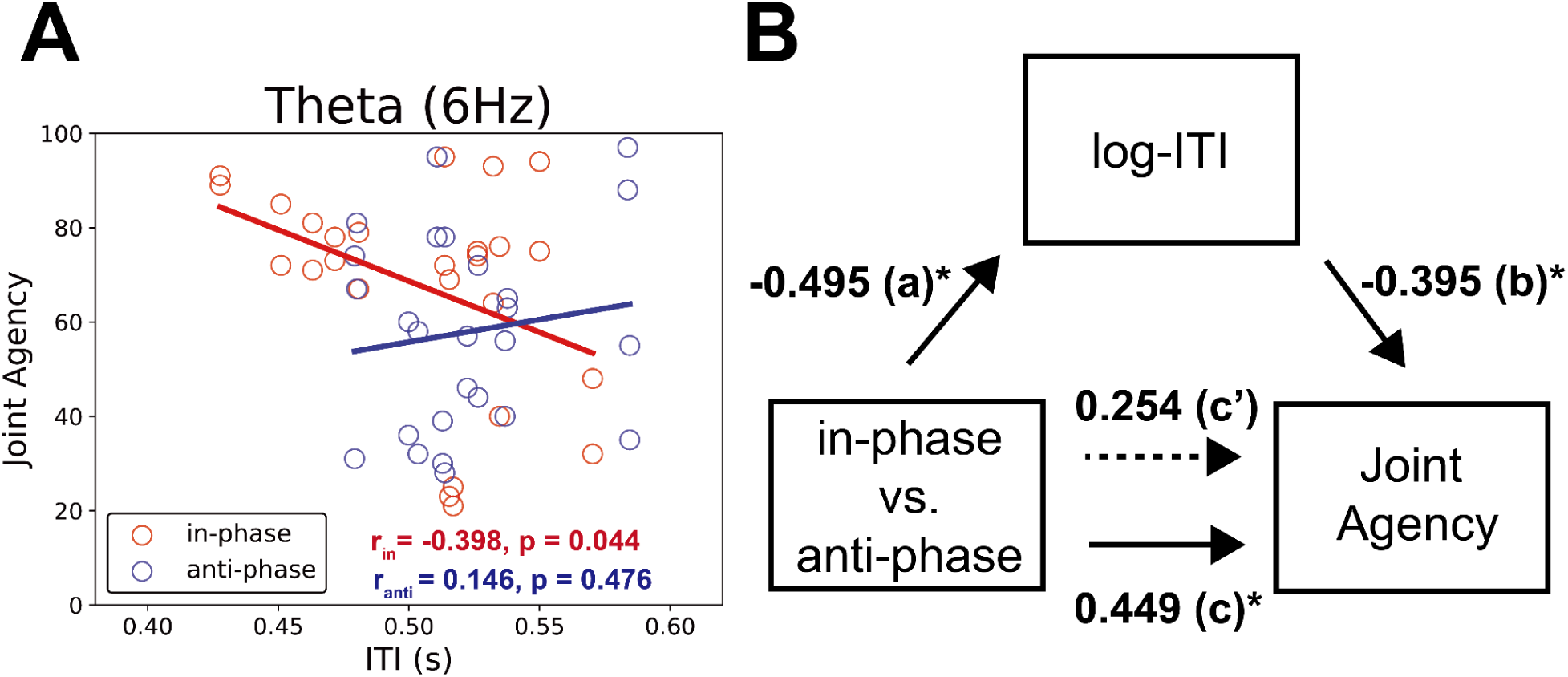
Correlation and mediation analyses between ITI and joint agency in the theta-frequency (6 Hz) stimulation. (A) In the in-phase mode, there was negative correlation between ITI and joint agency. (B) The effect of dual-stimulation on joint agency is partially mediated by log-transformed ITI (log-ITI). All path coefficients are standardized. *p < 0.05.

To test the hypothesis that tapping speed mediates the enhancement of joint agency, we conducted a mediation analysis (Figure 5B). In this model, path *a* represents the effect of stimulation phase on log-ITI, path *b* the association between log-ITI and joint agency controlling for phase, path *c* the total effect of phase on joint agency, and path *c′* the direct effect after accounting for log-ITI. The results indicated that (i) dual-brain stimulation (in-phase vs. anti-phase) could predict joint agency (path c = 0.449, p = 0.031) and that the relationship between dual-brain stimulation and joint agency was not significant when ITI was included in the model as a mediator (path c’ = 0.254, p = 0.208). The estimated indirect effect was positive (β = 0.196), with a 95% confidence interval that slightly crossed zero [−0.001, 0.670], suggesting a possible but not statistically robust mediation effect. In other words, ITI may partially mediate the effect of dual-brain stimulation on joint agency, but this indirect pathway should be interpreted cautiously given the uncertainty in the estimate and the modest sample size.

## Discussion

In the present study, we applied dual-tACS over the rTPJ while dyads performed an alternating tapping task to examine whether modulating putative inter-brain synchrony causally alters the sense of joint agency. Under theta (6 Hz) anti-phase stimulation, the tapping tempo was significantly slower than the memorized reference, whereas the other stimulation conditions did not show reliable deviations from the target tempo and relative-phase stability remained intact across all conditions. Subjectively, joint agency was lower in the theta anti-phase condition than in the theta in-phase condition, while IOS ratings were unaffected. Under theta stimulation, faster tapping predicted stronger joint agency only in the in-phase condition, and mediation analysis suggested that ITI partially accounted for the dual-tACS effect. These results indicate that disrupting theta-band alignment between participants’ rTPJ reduces the felt sense of acting together, in part by diminishing the efficiency of interpersonal coordination rather than by disrupting the structural alternation itself.

The present findings provide causal evidence that theta-band interpersonal alignment contributes to the subjective experience of joint agency, consistent with prior EEG hyperscanning work linking theta synchrony to interpersonal coordination. EEG hyperscanning reviews indicate that inter-brain theta coupling in frontal and parietal networks emerges when dyads co-attend and coordinate their actions ^13,14^. Consistent with this view, theta synchrony has been shown to increase with coordination demands ^21^. Léné et al. (2021) proposed that theta oscillations support a social-alignment feedback loop integrating mirror-neuron and predictive-coding mechanisms to dynamically couple perception and action between partners ^35^. In line with this theoretical framework, our phase-dependent modulation of joint agency by dual-tACS supports the idea that interpersonal theta-band alignment is closely involved in the construction of shared control.

The present findings also highlight regional specificity centered on the rTPJ. The rTPJ is widely implicated in mentalizing, perspective taking, and self–other distinction, core processes underlying how actions and intentions are attributed to oneself versus others ^28,29^. Hyperscanning studies repeatedly place the rTPJ within inter-brain networks engaged during joint action, with evidence showing coupling in this region scales with joint-agency ratings, coordination demands, and leadership roles ^16,31,36^. Converging causal evidence comes from neuromodulation studies showing that HD-tRNS over the rTPJ selectively alters attributions of humanness and cooperativeness toward a partner without affecting motor coordination ^37^. Extending this body of work, the present study suggests that dual-tACS targeting the rTPJ can modulate not only social judgments but also the subjective experience of acting together. By manipulating theta-phase alignment between partners, we altered joint agency while leaving the basic complementary coordination structure intact. Together, these findings support the view that the rTPJ contributes to joint agency as part of a broader theta-band interpersonal network supporting shared representations of action and agency, rather than serving as a standalone generator of coordinated behavior.

The behavioral data indicate that theta anti-phase stimulation selectively reduced the efficiency with which dyads maintained the target tempo while leaving the stability of complementary coordination unchanged. Under this condition, dyads exhibited a slower tapping tempo, whereas the 180° alternation and phase variability (SDφ) remained comparable across conditions. This dissociation suggests that rTPJ theta misalignment does not impair the ability to sustain complementary roles per se, but instead makes it harder for partners to jointly implement a memorized tempo efficiently. One plausible interpretation is that anti-phase stimulation increased the neural demands associated with online prediction of the partner’s next tap, leading participants to adopt a more conservative, slower pace to preserve coordination. From the participants’ perspective, such compensatory slowing may have been experienced as less fluent or less “shared,” which could explain the reduction in subjective joint agency despite the preserved coordination structure. In line with this idea, noninvasive brain stimulation has been shown to alter subjective effort without markedly changing motor performance; for example, cathodal transcranial direct current stimulation (tDCS) over the left dorsolateral prefrontal cortex kept leg-press volume load comparable to sham stimulation, while significantly increasing ratings of perceived exertion ^38^. Our pattern of reduced tempo efficiency and diminished joint agency, alongside an intact coordination structure, may reflect a similar “unchanged performance but increased internal effort” type of effect at the level of interpersonal coordination.

Our mediation analysis is consistent with the view that the dual-tACS effect on joint agency may be partly transmitted through changes in coordination efficiency. Under theta in-phase stimulation, faster tapping (i.e., shorter inter-tap intervals) was associated with stronger joint agency, whereas no reliable relationship between tempo and joint agency emerged under theta anti-phase stimulation. When the phase condition and inter-tap interval were included in the same model, inter-tap interval statistically accounted for part of the phase effect on joint agency, although this indirect pathway should be interpreted cautiously given the modest sample size and the wide confidence interval around the mediation estimate. This pattern echoes the mediation reported by Pan et al. (2021), where 6-Hz in-phase dual-tACS over frontal regions improved intonation learning in part via increased spontaneous movement synchrony between partners ^19^. Taken together, these findings suggest that theta-specific, phase-sensitive dual-tACS can influence higher-order subjective outcomes, such as the sense of joint agency, partly through its impact on coordination.

Joint agency and IOS showed a clear dissociation in our data: dual-tACS modulated perceived shared control but did not alter perceived interpersonal closeness. This dissociation may reflect a distinction between moment-to-moment shared control over an action and more enduring relational closeness ^39^. In the present study, dyads were acquaintances or friends who already reported relatively high IOS scores, so the stimulation may have been too subtle or too brief to affect this more stable aspect of the relationship. The dissociation instead points to a more specific role of rTPJ-centered theta processes in supporting shared control during joint action, rather than in shaping general interpersonal affinity.

Several limitations should be noted. First, our sample size of 13 dyads (26 individuals) is smaller than in many dual-brain modulation studies on interpersonal coordination and social learning, which often test around 20–30 dyads ^18,19,40^. Within this dataset, we nonetheless observed clear theta-specific, phase-dependent differences in tapping tempo and joint agency, but the limited N reduces the precision of the estimated effects and may obscure smaller or more nuanced patterns. Replication in larger and more diverse samples will therefore be important for assessing the robustness and boundary conditions of the present findings. Second, we stimulated only the rTPJ and did not include a sham condition. Although the frequency- and phase-specific pattern of results argues against a purely nonspecific effect, the contribution of peripheral sensations or general arousal cannot be fully ruled out; future work should incorporate double-blind sham and montage-control conditions to strengthen causal inferences. Moreover, previous hyperscanning work has shown that theta-band inter-brain synchrony between prefrontal regions and the rTPJ scales with joint agency ratings during alternating tapping ^16^, suggesting that stimulating additional nodes of this network (e.g., prefrontal sites or multi-site montages) may be necessary to fully characterize the causal contributions of the broader rTPJ-centered circuit. Third, we inferred changes in inter-brain synchrony from the stimulation protocol rather than measuring neural coupling directly. Dual-tACS is designed to modulate inter-brain phase relations, but the actual degree of entrainment at the cortical level remains unknown. Combining dual-tACS with EEG hyperscanning would allow direct tests of whether in-phase stimulation enhances rTPJ theta coupling and whether such changes track moment-to-moment fluctuations in joint agency ^41^. Fourth, the study focused on same-gender dyads who were already acquainted. While this choice reduces variability due to gender composition and unfamiliarity, it limits generalizability to mixed-gender or stranger dyads, where social dynamics and the baseline sense of agency may differ. Finally, we examined only one type of joint action—alternating rhythmic tapping with a fixed target tempo. In more complex cooperative contexts, such as musical performance or joint problem solving, the relationship between inter-brain synchrony, coordination, and joint agency may involve additional cognitive and affective factors. These limitations highlight the need to examine broader task contexts and to adopt converging methodological approaches in future research.

While the present study relied on transcranial electrical stimulation, future studies could use other neuromodulatory methods. Recent work suggests that sensory multi-brain stimulation, such as synchronized visual flicker, can enhance dyadic cooperation by entraining neural oscillations and increasing inter-brain synchrony ^40^. Comparing electrical and sensory multi-brain stimulation—or combining them—may help clarify whether similar changes in joint agency can be achieved through distinct neural routes to inter-brain alignment.

In this study, we used dual-tACS over the rTPJ while dyads performed an alternating tapping task to test whether manipulating theta-band phase alignment between partners’ brains alters the sense of joint agency. Theta anti-phase stimulation slowed the tapping tempo relative to the memorized reference and reduced perceived joint agency compared with theta in-phase stimulation, while the 180° alternation structure of coordination remained intact. Inter-tap interval may partially mediate this phase effect, although this indirect pathway should be interpreted cautiously. Ratings of interpersonal closeness, assessed with the IOS scale, were unchanged, indicating that rTPJ-centered theta alignment is more closely related to moment-to-moment shared control than to broader relationship closeness. Together, these findings suggest that theta-band interpersonal alignment in networks involving the rTPJ shapes how efficiently partners coordinate and how strongly they experience their actions as jointly controlled.

## Resource availability

### Lead contact

Requests for further information and resources should be directed to, and will be fulfilled by, the lead contact,Yuto Kurihara

### Materials availability

This study has not generated any new reagents.

### Data and code availability

The tapping data and analysis code will be deposited in OSF and made publicly available upon publication. The OSF URL/DOI will be provided in the final version of the manuscript. De-identified questionnaire data will be available from the lead contact upon reasonable request, subject to institutional ethics and participant privacy restrictions. Any additional information required to reanalyze the data reported in this paper will be available from the lead contact upon request.

## Acknowledge

We thank Akito Miura for providing the facilities and equipment necessary to conduct the dual-brain stimulation experiment. We also thank Vincent Chamberland for helpful comments and careful editing of the manuscript. This study was supported by the Japan Society for the Promotion of Science (JSPS) KAKENHI Grant-in-Aid for Early-Career Scientists (24K21066 to Y.K.) and Grant-in-Aid for Scientific Research (A) (21H04425 to R.O.). G.D. was supported by the Institute for Data Valorization, Montreal and the Canada First Research Excellence Fund (IVADO; CF00137433), the Fonds de recherche du Québec (FRQ; 285289), the Natural Sciences and Engineering Research Council of Canada (NSERC; DGECR-2023-00089), and the Canadian Institute for Health Research (CIHR 192031; SCALE).

## Author contributions

Y.K. and A.T. designed this research. Y.K. analyzed acquired data. Y.K. prepared figures; Y.K. drafted manuscript; Y.K., G.D., and R.O. edited and revised manuscript; Y.K., A.Y., G.D., and R.O. approved final version of manuscript.

## Declaration of generative AI and AI-assisted technologies in the writing process

During the preparation of this work the authors used ChatGPT in order to improve the language of the manuscript. After using this tool, the authors reviewed and edited the content as needed and took full responsibility for the content of the published article.

## STAR Methods

## Key resources table

| REAGENT or RESOURCE | SOURCE | IDENTIFIER |
| --- | --- | --- |
| Deposited data |  |  |
| Tapping dataset | This paper | OSF: to be provided upon acceptance |
| Analysis code | This paper | OSF: to be provided upon acceptance |
| Software and algorithms |  |  |
| MATLAB | MathWorks | MATLAB R2025a |
| COMETS Toolbox | Saturnino et al. | version 2.0 |
| Python | Python Software Foundation | version 3.13.5 |
| SciPy | SciPy community | version 1.15.3 |
| pingouin | pingouin developers | version 0.5.5 |
| Rstudio | Posit Software, PBC | 2025.09.1+401 |
| lme4 | Bates et al. / CRAN | version 1.1-37 |
| lmerTest | Kuznetsova et al. / CRAN | version 3.1-3 |
| **Other resources** |  |  |
| Nurostym tACS device | Neuro Device Group S.A.,<br>Warszawa, Poland | https://brainbox-neuro.com/products/nurostym-tes |
| Inclusion of Other in the Self (IOS) scale | Aron et al. | cited in main text |
| Joint agency visual analog scale | This paper | see STAR Methods |
| tES side-effect questionnaire | This paper / adapted from prior tES side-effect scales | see STAR Methods |

## Method details

### Participants

Thirteen same-gender dyads were recruited for this study (8 female pairs; mean age =22.48 years, SD =3.05, range = 18–30). All dyads were acquainted prior to enrollment. All participants provided written informed consent. The experimental procedures adhered to the Declaration of Helsinki’s code of ethics for human experimentation and received approval from the Waseda University Ethical Review Committee.

### Alternative tapping task

Participants in each dyad were seated side by side and were asked not to look at their partner’s face or fingers. Participants were approximately 70 cm apart and instructed to coordinate alternating (anti-phase) tapping with two computer mice. Participants used their right index finger to click the left button of a computer mouse, which was placed on one of two 40 × 50 cm tables. Each tap produced an auditory feedback sound at a frequency of 440 Hz. In the tapping task, participants initially listened to a reference tempo (tapping speed = 0.5s). Following the transmission of the first eight sounds, the metronome was deactivated. Subsequently, the pairs began tapping at a tempo as close as possible to the memorized reference tempo. Each session was completed after 300 taps. The mean time required to complete the tapping task in each session and condition was 156.00 s (SD = 2.30 s).

### Dual-transcranial alternating current stimulation system

During the alternating tapping, two Nurostym tACS devices (Neuro Device Group S.A., Warszawa, Poland) simultaneously delivered sinusoidal electrical current signals to the brains of both participants. Transcranial signals were delivered via two sponge electrodes (anodal: 3 × 4 cm, cathodal: 5 × 7 cm) that were soaked in a saline solution. The smaller anodal electrode was placed over the rTPJ (equivalent to electrode position CP6 according to the international 10/10 system), and the larger cathodal electrode on the contralateral (left) forearm. For tACS, the current intensity was set at 3.0 mA. This intensity falls within the range classified as low-intensity transcranial electric stimulation (tES), for which large-scale safety reviews and guidelines report no serious adverse events when the current remains below 4 mA and the total stimulation duration does not exceed approximately 60 min per day ^42,43^. In our protocol, stimulation blocks were relatively brief and the total stimulation duration per participant was well below these recommended limits. Post-session questionnaires confirmed only mild, transient sensations (e.g., tingling, itching) with no serious or lasting adverse effects. Using the COMETS Toolbox (version 2.0) ^44^, we simulated the induced electric field and confirmed that this montage produced a maximal field over the right temporo-parietal junction (around CP6), indicating that it is suitable for entraining neural activity in this region (Figure 1B). The two stimulators were controlled in MATLAB R2025a (MathWorks, Natick, MA, USA) using Data Acquisition Toolbox and the Data Acquisition Toolbox Support Package for National Instruments NI-DAQmx Devices. The tACS used sinusoidal stimulation at 6, 10, and 20 Hz, delivered in-phase or anti-phase between dyad participants.

### Procedure

Upon arrival, participants were briefed on the experimental procedures, and tACS electrodes were positioned. The experiments consisted of six sessions conducted under six tACS coupling conditions, defined by the factorial combination of three stimulation frequencies (6, 10, 20 Hz) and two inter-dyad phase relationships (in-phase, anti-phase). To achieve counterbalancing, the six sessions were randomized within each dyad. Which member of the dyad acted as the first and second mover (i.e., who initiated tapping) was fixed across all sessions. We asked each participant to rate their joint agency using a visual analog scale (VAS) at the end of each session. The instructional text of joint agency is as follows: *During the tapping exercise, to what extent did you feel that you were jointly controlling the rhythm and tempo with your partner? Please indicate your response on the scale, where the rightmost end represents ‘I was jointly controlling it to the greatest extent,’ and the leftmost end represents ‘I was not jointly controlling it at all (I was controlling it independently)*.’ After rating their sense of joint agency at the end of each session, participants completed the Inclusion of Other in the Self (IOS) scale ^34^ to assess their perceived closeness to their partner during that session. After finishing the six sessions, participants completed a questionnaire assessing tES-related side effects. They rated the intensity of several sensations (tingling/itching, pain, burning sensation, warmth/heat, metallic/iron taste, fatigue/drowsiness, and any other sensations) on a four-point scale (0 = none, 1 = mild, 2 = moderate, 3 = severe).

## Quantification and statistical analysis

### Behavioral analysis

To calculate the inter-tap interval (ITI) in the combined tap sequence of participants A and B, we subtracted the tap onset time from the subsequent partner’s tap onset time:

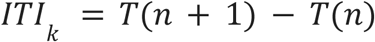

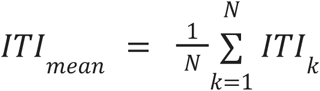

where T(n) denotes the time of the n-th tap in the combined tap sequence, which may be produced by either participant A or participant B. The ITI_mean_ is the average of ITI.

We also calculated relative phase (RP) of alternating tapping using a circular measure to confirm whether their tapping was clustered around anti-phase (RP = 180 deg.). RP was defined as follows:

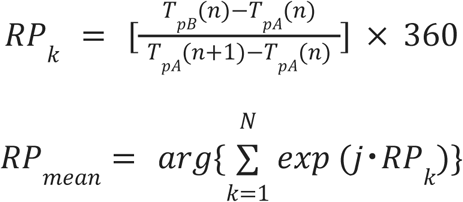

where TpA (n) and TpB (n) stand for the n-th tapping time of participants A and B, respectively. The RP_mean_ is the average of RP which is calculated by circular statistics. The j shows the complex number and the arg{・} shows the argument of complexity. The RP range was 0 to 360 deg. In addition, we calculated the standard deviation of relative phase (SDφ) to evaluate the stability of alternative tapping. The SDφ were calculated by circular statistics as below:

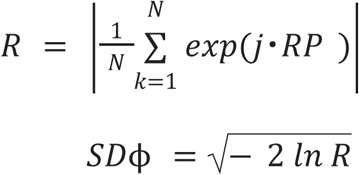

where SDφ represents the instability of alternative tapping. R is the resultant length of the RP vectors.

### Statistical analysis

To assess adherence to the reference tempo (0.50 s), we conducted one-sample t-tests against μ = 0.50 s for each of the six conditions (Frequency: 6, 10, 20 Hz × Phase: in-phase, anti-phase), with p-values FDR-corrected across the six tests. Similarly, to verify that tapping was performed at the target relative phase of 180°, we performed one-sample t-tests against μ = 180° for each condition, with p-values FDR-corrected across the six tests (3 frequencies × 2 phases). For tapping stability, we conducted a two-way repeated-measures ANOVA with within-subject factors Frequency (6, 10, 20 Hz) and Phase (in-phase, anti-phase). For the subjective measures, we conducted two-way ANOVAs with within-subject factors Frequency (6, 10, 20 Hz) and Phase (in-phase, anti-phase) on joint agency and on the IOS scale. One-sample t-tests were performed using SciPy (version 1.15.3), and two-way ANOVAs were conducted with *pingouin* (version 0.5.5), both in a Python 3.13.5 environment.

We performed correlation and mediation analyses to explore potential relationships between distinct dependent variables ^19,45^: ITI and joint agency. Following up on the results from the previous analyses of tapping and subjective scores, we defined the stimulation mode as the independent variable, ITI as the mediator, and joint agency as the dependent variable. The mediation model was defined as follows:

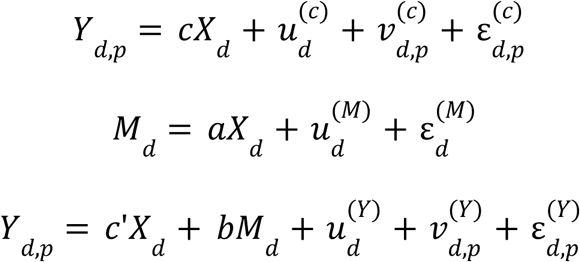

where d is the dyad number (ID), and p is individual participant ID. X is the independent variable (phase mode of dual-brain stimulation, dummy coded, with 1 for in-phase and 0 for anti-phase), M is the mediator (log-transformed inter-tap interval; log-ITI), Y is the dependent variable (Joint agency), u and v are the random intercepts of dyads or individual persons, and ε denotes the residuals. Path a is the coefficient relating X to M. Path _b_ is the coefficient relating M to Y adjusted for X. Paths c’ and c describe the relationship of Y and X with and without M, respectively. We used bootstrapping procedures implemented in Rstudio (version 2025.09.1+401), using the lme4 (version 1.1.37) and lmerTest (version 3.1.3) packages, to determine whether the mediation effect was different from zero with 95% confidence. The number of bootstrap samples was set to 5000.

## References

1. Gallagher, S. (2000). Philosophical conceptions of the self: implications for cognitive science. Trends Cogn. Sci. 4, 14–21. 10.1016/S1364-6613(99)01417-5.

2. Haggard, P., and Tsakiris, M. (2009). The Experience of Agency: Feelings, Judgments, and Responsibility. Curr. Dir. Psychol. Sci. 18, 242–246. 10.1111/j.1467-8721.2009.01644.x.

3. David, N., Newen, A., and Vogeley, K. (2008). The “sense of agency” and its underlying cognitive and neural mechanisms. Conscious. Cogn. 17, 523–534. 10.1016/j.concog.2008.03.004.

4. Decety, J., and Lamm, C. (2007). The Role of the Right Temporoparietal Junction in Social Interaction: How Low-Level Computational Processes Contribute to Meta-Cognition. The Neuroscientist 13, 580–593. 10.1177/1073858407304654.

5. Haggard, P., and Chambon, V. (2012). Sense of agency. Curr. Biol. 22, R390–R392. 10.1016/j.cub.2012.02.040.

6. Haggard, P. (2017). Sense of agency in the human brain. Nat. Rev. Neurosci. 18, 196–207. 10.1038/nrn.2017.14.

7. Sebanz, N., Bekkering, H., and Knoblich, G. (2006). Joint action: Bodies and minds moving together. Trends Cogn. Sci. 10, 70–76. 10.1016/j.tics.2005.12.009.

8. Pacherie, E. (2012). The Phenomenology of Joint Action: Self-Agency vs. Joint-Agency. In Joint Attention: New Developments, A. Seemann, ed. (MIT Press), pp. 343–389.

9. Zapparoli, L., Paulesu, E., Mariano, M., Ravani, A., and Sacheli, L.M. (2022). The sense of agency in joint actions: A theory-driven meta-analysis. Cortex 148, 99–120. 10.1016/j.cortex.2022.01.002.

10. Loehr, J.D. (2022). The sense of agency in joint action: An integrative review. Psychon. Bull. Rev. 29, 1089–1117. 10.3758/s13423-021-02051-3.

11. Bolt, N.K., Poncelet, E.M., Schultz, B.G., and Loehr, J.D. (2016). Mutual coordination strengthens the sense of joint agency in cooperative joint action. Conscious. Cogn. 46, 173–187. 10.1016/j.concog.2016.10.001.

12. Zhou, Z., Christensen, J., Cummings, J.A., and Loehr, J.D. (2023). Not just in sync: Relations between partners’ actions influence the sense of joint agency during joint action. Conscious. Cogn. 111, 103521. 10.1016/j.concog.2023.103521.

13. Czeszumski, A., Eustergerling, S., Lang, A., Menrath, D., Gerstenberger, M., Schuberth, S., Schreiber, F., Rendon, Z.Z., and König, P. (2020). Hyperscanning: A Valid Method to Study Neural Inter-brain Underpinnings of Social Interaction. Front. Hum. Neurosci. 14. 10.3389/fnhum.2020.00039.

14. De Fano, A., Fiedler, P., Zappasodi, F., Bertollo, M., and Comani, S. (2025). Non-verbal joint action in healthy adults: a systematic scoping review of EEG-hyperscanning research. Soc. Cogn. Affect. Neurosci. 20, nsaf050. 10.1093/scan/nsaf050.

15. Dumas, G., Nadel, J., Soussignan, R., Martinerie, J., and Garnero, L. (2010). Inter-Brain Synchronization during Social Interaction. PLOS ONE 5, e12166. 10.1371/journal.pone.0012166.

16. Shiraishi, M., and Shimada, S. (2021). Inter-brain synchronization during a cooperative task reflects the sense of joint agency. Neuropsychologia 154, 107770. 10.1016/j.neuropsychologia.2021.107770.

17. Lu, H., Wang, X., Zhang, Y., Huang, P., Xing, C., Zhang, M., and Zhu, X. (2023). Increased interbrain synchronization and neural efficiency of the frontal cortex to enhance human coordinative behavior: A combined hyper-tES and fNIRS study. NeuroImage 282, 120385. 10.1016/j.neuroimage.2023.120385.

18. Novembre, G., Knoblich, G., Dunne, L., and Keller, P.E. (2017). Interpersonal synchrony enhanced through 20 Hz phase-coupled dual brain stimulation. Soc. Cogn. Affect. Neurosci. 12, 662–670. 10.1093/scan/nsw172.

19. Pan, Y., Novembre, G., Song, B., Zhu, Y., and Hu, Y. (2021). Dual brain stimulation enhances interpersonal learning through spontaneous movement synchrony. Soc. Cogn. Affect. Neurosci. 16, 210–221. 10.1093/scan/nsaa080.

20. Szymanski, C., Müller, V., Brick, T.R., von Oertzen, T., and Lindenberger, U. (2017). Hyper-Transcranial Alternating Current Stimulation: Experimental Manipulation of Inter-Brain Synchrony. Front. Hum. Neurosci. 11. 10.3389/fnhum.2017.00539.

21. Kurihara, Y., Takahashi, T., and Osu, R. (2022). The relationship between stability of interpersonal coordination and inter-brain EEG synchronization during anti-phase tapping. Sci. Rep. 12. 10.1038/s41598-022-10049-7.

22. Kurihara, Y., Takahashi, T., and Osu, R. (2024). The topology of interpersonal neural network in weak social ties. Sci. Rep. 14, 4961. 10.1038/s41598-024-55495-7.

23. Novembre, G., and Iannetti, G.D. (2021). Hyperscanning Alone Cannot Prove Causality. Multibrain Stimulation Can. Trends Cogn. Sci. 25, 96–99. 10.1016/j.tics.2020.11.003.

24. Helfrich, R.F., Schneider, T.R., Rach, S., Trautmann-Lengsfeld, S.A., Engel, A.K., and Herrmann, C.S. (2014). Entrainment of Brain Oscillations by Transcranial Alternating Current Stimulation. Curr. Biol. 24, 333–339. 10.1016/j.cub.2013.12.041.

25. Feurra, M., Bianco, G., Santarnecchi, E., Del Testa, M., Rossi, A., and Rossi, S. (2011). Frequency-dependent tuning of the human motor system induced by transcranial oscillatory potentials. J. Neurosci. Off. J. Soc. Neurosci. 31, 12165–12170. 10.1523/JNEUROSCI.0978-11.2011.

26. Polanía, R., Nitsche, M.A., Korman, C., Batsikadze, G., and Paulus, W. (2012). The Importance of Timing in Segregated Theta Phase-Coupling for Cognitive Performance. Curr. Biol. 22, 1314–1318. 10.1016/j.cub.2012.05.021.

27. Carter, R.M., and Huettel, S.A. (2013). A nexus model of the temporal–parietal junction. Trends Cogn. Sci. 17, 328–336. 10.1016/j.tics.2013.05.007.

28. Gallagher, H.L., and Frith, C.D. (2003). Functional imaging of ‘theory of mind.’ Trends Cogn. Sci. 7, 77–83. 10.1016/S1364-6613(02)00025-6.

29. Saxe, R., and Kanwisher, N. (2003). People thinking about thinking people: The role of the temporo-parietal junction in “theory of mind.” NeuroImage 19, 1835–1842. 10.1016/s1053-8119(03)00230-1.

30. Jahng, J., Kralik, J.D., Hwang, D.-U., and Jeong, J. (2017). Neural dynamics of two players when using nonverbal cues to gauge intentions to cooperate during the Prisoner’s Dilemma Game. NeuroImage 157, 263–274. 10.1016/j.neuroimage.2017.06.024.

31. Jiang, J., Chen, C., Dai, B., Shi, G., Ding, G., Liu, L., and Lu, C. (2015). Leader emergence through interpersonal neural synchronization. Proc. Natl. Acad. Sci. 112, 4274–4279. 10.1073/pnas.1422930112.

32. Tang, H., Mai, X., Wang, S., Zhu, C., Krueger, F., and Liu, C. (2016). Interpersonal brain synchronization in the right temporo-parietal junction during face-to-face economic exchange. Soc. Cogn. Affect. Neurosci. 11, 23–32. 10.1093/scan/nsv092.

33. Wang, L., Chang, Y., Liou, S., Weng, M., Chen, D., and Kung, C. (2024). When “more for others, less for self” leads to co-benefits: A tri-MRI dyad-hyperscanning study. Psychophysiology 61. 10.1111/psyp.14560.

34. Aron, A., Aron, E.N., and Smollan, D. (1992). Inclusion of Other in the Self Scale and the structure of interpersonal closeness. J. Pers. Soc. Psychol. 63, 596–612. 10.1037/0022-3514.63.4.596.

35. Léné, P., Karran, A.J., Labonté-Lemoyne, E., Sénécal, S., Fredette, M., Johnson, K.J., and Léger, P.-M. (2021). Is there collaboration specific neurophysiological activation during collaborative task activity? An analysis of brain responses using electroencephalography and hyperscanning. Brain Behav. 11, e2270. 10.1002/brb3.2270.

36. Abe, M.O., Koike, T., Okazaki, S., Sugawara, S.K., Takahashi, K., Watanabe, K., and Sadato, N. (2019). Neural correlates of online cooperation during joint force production. NeuroImage 191, 150–161. 10.1016/j.neuroimage.2019.02.003.

37. Moreau, Q., Chamberland, V., Moses, L., Milanova, G., and Dumas, G. (2025). Online HD-tRNS Over the Right Temporoparietal Junction Modulates Social Inference But Not Motor Coordination. eNeuro 12. 10.1523/ENEURO.0155-25.2025.

38. Lattari, E., Rosa Filho, B.J., Fonseca Junior, S.J., Murillo-Rodriguez, E., Rocha, N., Machado, S., and Maranhão Neto, G.A. (2020). Effects on Volume Load and Ratings of Perceived Exertion in Individuals’ Advanced Weight Training After Transcranial Direct Current Stimulation. J. Strength Cond. Res. 34, 89. 10.1519/JSC.0000000000002434.

39. Liepelt, R., and Raab, M. (2021). Metacontrol and joint action: how shared goals transfer from one task to another? Psychol. Res. 85, 2769–2781. 10.1007/s00426-020-01443-9.

40. Leiva-Cisterna, I., Barraza, P., Rodríguez, E., and Dumas, G. (2025). Sensory multi-brain stimulation enhances dyadic cooperative behavior. Soc. Cogn. Affect. Neurosci. 20, nsaf104. 10.1093/scan/nsaf104.

41. Takeuchi, N. (2024). A dual-brain therapeutic approach using noninvasive brain stimulation based on two-person neuroscience: A perspective review. Math. Biosci. Eng. MBE 21, 5118–5137. 10.3934/mbe.2024226.

42. Antal, A., Alekseichuk, I., Bikson, M., Brockmöller, J., Brunoni, A.R., Chen, R., Cohen, L.G., Dowthwaite, G., Ellrich, J., Flöel, A., et al. (2017). Low intensity transcranial electric stimulation: Safety, ethical, legal regulatory and application guidelines. Clin. Neurophysiol. 128, 1774–1809. 10.1016/j.clinph.2017.06.001.

43. Matsumoto, H., and Ugawa, Y. (2017). Adverse events of tDCS and tACS: A review. Clin. Neurophysiol. Pract. 2, 19–25. 10.1016/j.cnp.2016.12.003.

44. Lee, C., Jung, Y.-J., Lee, S.J., and Im, C.-H. (2017). COMETS2: An advanced MATLAB toolbox for the numerical analysis of electric fields generated by transcranial direct current stimulation. J. Neurosci. Methods 277, 56–62. 10.1016/j.jneumeth.2016.12.008.

45. Preacher, K.J., and Hayes, A.F. (2008). Asymptotic and resampling strategies for assessing and comparing indirect effects in multiple mediator models. Behav. Res. Methods 40, 879–891. 10.3758/BRM.40.3.879.

